# Contextual and spatial associations between objects interactively modulate visual processing

**DOI:** 10.1101/2020.05.20.106070

**Authors:** Genevieve L. Quek, Marius V. Peelen

## Abstract

Much of what we know about object recognition arises from the study of isolated objects. In the real world, however, we commonly encounter groups of contextually-associated objects (e.g., teacup, saucer), often in stereotypical spatial configurations (e.g., teacup *above* saucer). Here we used EEG to test whether identity-based associations between objects (e.g., teacup-saucer *vs*. teacup-stapler) are encoded jointly with their typical relative positioning (e.g., teacup *above* saucer *vs. below* saucer). Observers viewed a 2.5Hz image stream of contextually-associated object pairs intermixed with non-associated pairs as every fourth image. The differential response to non-associated pairs (measurable at 0.625Hz in 28/37 participants), served as an index of contextual integration, reflecting the association of object identities in each pair. Over right occipitotemporal sites, this signal was larger for typically-positioned object streams, indicating that spatial configuration facilitated the extraction of the objects’ contextual association. This high-level influence of spatial configuration on object identity integration arose ∼320ms post stimulus onset, with lower-level perceptual grouping (shared with inverted displays) present at ∼130ms. These results demonstrate that contextual and spatial associations between objects interactively influence object processing. We interpret these findings as reflecting the high-level perceptual grouping of objects that frequently co-occur in highly stereotyped relative positions.

---

Real-world visual environments are characterised by strong contextual associations between objects. These identity-based associations have been a focus of investigation for nearly half a century, and there is now overwhelming evidence to suggest that the co-occurrence statistics of objects (i.e., the likelihood that two objects are found in the same environment) influence observers’ ability to recognise and remember these objects (Biederman 1972; Biederman et al. 1973; Hock et al. 1974; Palmer 1975; Biederman et al. 1982; Holcomb and Mcpherson 1994; Ganis and Kutas 2003; Davenport and Potter 2004; Davenport 2007; Munneke et al. 2013).

In addition to identity-based contextual associations, objects in real-world scenes also occupy highly regular positions, both in an absolute sense (e.g., airplanes appear in the sky) but also relative to each other (lamps are found above tables, not below tables). There is evidence that such positional regularities between objects also modulate object processing (Bar and Ullman 1996; Riddoch et al. 2003; Green and Hummel 2006; Gronau and Shachar 2014), with recent studies reporting that the spatial configuration of real-world objects impacts the way observers search for, represent, and perceive these stimuli (e.g, Boettcher et al. 2018; for recent reviews, see Kaiser et al. 2019; Võ et al. 2019).

In the present EEG study, we asked how identity-based and positional regularities interact during visual object processing. We hypothesized that identity-based associations between objects are encoded jointly with the objects’ typical relative position, with both types of regularities needed for creating a coherent group (Bar and Ullman 1996). For example, a lamp above a table creates a coherent and familiar group, with this coherence being disrupted by changing either the identity or the position of one of the objects. Therefore, we predicted that neural responses indexing identity-based associations (e.g., lamp-table *vs*. lamp-bush) should depend on the spatial configuration of the associated objects being correct. Furthermore, in line with interactive models of object and scene perception (Bar 2004; Davenport and Potter 2004; Brandman and Peelen 2017), we predicted that this combination of identity-based and positional regularity would modulate object representations during perceptual processing stages, that is, in high-level visual cortex.

An alternative possibility is that identity-based contextual associations are independent of positional regularity, with the contextually-associated objects processed and recognized independently of each other, before being associated at later processing stages. This would be in line with the classical view of the visual system as creating positionally-invariant object representations. Namely, if individual object identity is first accessible at a position-invariant stage of visual processing, the relative positioning of real world objects might have little impact on identity-based associations, since by the time the objects’ identity can be ‘read out’ and compared, information about spatial location is largely discarded.

While some studies have provided neural evidence for interactions between identity-based and positional regularities during object perception (Gronau et al. 2008; Baeck et al. 2013), others have shown no such interaction, reporting similar effects of positional regularity (e.g., being correctly positioned for an action) for contextually-associated and non-associated objects in object selective cortex (Roberts and Humphreys 2010; Kim and Biederman 2011).

There are two important issues that should be considered when interpreting these results: First, in the studies reporting interactions between identity-based and positional regularities, the relationship between the objects was task-relevant, such that participants likely used cognitive strategies (e.g., differential attentional allocation) based on their knowledge of the relationship between the displayed objects. For example, in one fMRI study, differences between regularly and irregularly positioned action pairs in object-selective cortex were evident only when participants judged the correctness of the configuration of the objects, and not when they performed an orthogonal task (Baeck *et al*. 2013). Still other studies have explicitly focused on behavioural benefits of identity-based and positional regularities, for example in sequential displays (Gronau *et al*. 2008), necessarily making these regularities task-relevant. This leaves open the question of whether identity-based and positional regularities affect neural processing in the absence of an explicit task involving these regularities.

Second, contextually-associated and regularly-positioned objects in the real world often share low-level visual features and are typically aligned, such that they can be grouped based on Gestalt principles (Wagemans et al. 2012). For example, correctly positioned objects such as a screwdriver and a screw (e.g., Green and Hummel 2006) share visual features (elongation) and are grouped based on their alignment. The lack of a visual control condition in previous studies leaves open the possibility that these results partly reflected differences in low-level feature similarity and/or Gestalt-level grouping between conditions.

In the present electroencephalography (EEG) study, we take advantage of the steady-state visual evoked potential (Regan 1966; Regan 1989; Norcia et al. 2015) that is measurable on the scalp when a visual stimulus is presented at a strict periodicity (Tononi et al. 1998; Baldauf and Desimone 2014). The logic behind the specific frequency-tagging variation we use here has been described elsewhere (Liu-Shuang et al. 2014; Rossion et al. 2015); here we exploited this approach to test for the interaction between identity-based and positional object associations that were task-irrelevant, while controlling for possible low-level confounds. Observers viewed a rapid stream (2.5 Hz presentation rate) of contextually-associated object pairs (i.e., ‘match’ pairs) intermixed with pairs of non-associated objects (i.e., ‘mismatch’ pairs, presented as every fourth stimulus for a mismatch frequency of 0.625 Hz) (see movie in Supplemental Material). The selective response to these mismatch pairs provides a highly sensitive index of the degree to which identity-based associations are processed in the absence of an explicit task. In this way, the differential response to non-associated pairs presented among contextually-associated pairs acts as a proxy for *contextual integration*, reflecting the association of object identities in each pair. Here we refer to this identity-based association index interchangeably as the ‘mismatch-selective response’ and ‘contextual integration response’.

The central manipulation in our study was the relative position of the two objects making up the pairs in the stream: these were positioned either typically (e.g., lamp above table) or atypically (lamp below table), while everything else remained identical. We thus asked whether the Contextual Integration Response (i.e., the response at the mismatch frequency) was sensitive to the relative positional regularity of the two objects. In contrast to previous studies, we did not direct participants’ attention to the *associations* between objects, but rather engaged them in an orthogonal target-detection task that rendered both the identity-based and positional regularities in the pairs task-irrelevant. Here, participants monitored the sequence for images of handbags or telephones (which never appeared at the mismatch frequency), enabling us to capture a task-independent neural index of identity-based object associations. Crucially, we also presented all stimulus conditions upside-down (i.e., display rotated by 180 degrees). This inversion manipulation preserves all low-level properties (e.g., the featural similarity and alignment of the objects) while disrupting higher-level integration. Together, these design features allowed us to test for the interaction between identity-based and position-based associations between objects during visual processing.

To anticipate our results, we find that the degree to which the visual system integrates identity information carried by concurrent objects critically depends on their relative position, an effect that was most pronounced over right occipitotemporal cortex. This effect was eliminated by inverting the display, ruling out the possibility that the enhanced identity-based integration signal for typical configurations reflected low-level (Gestalt-like) grouping. Taken together, our results provide evidence that the spatial and identity relations between objects influence object processing in an interactive manner.

## Materials & Methods

### Participants

37 English-speaking individuals (17 males) aged between 19-34 years (*M =* 23.7 ± 4.1 years) took part in the experiment in exchange for monetary compensation (33 right handed, 3 left handed, 1 unreported). All reported having normal or corrected-to-normal vision and no neurological or psychiatric history. Prior to experimental testing all participants gave their written informed consent. Experimental data were stored under pseudonymised codes in accordance with the European General Data Protection Regulation (EU GDPR). The study was approved by the Radboud University Faculty of Social Sciences Ethics Committee (ECSW2017-2306-517).

### Stimuli

Stimuli were 48 individual object images (Kaiser et al. 2014) taken from 12 distinct real world contexts (bathroom, breakfast, closet, kitchen, dining, stove, beach, office, tea, bar, toilet, film). For each context (e.g., bathroom) there were two exemplars of a ‘top’ object (e.g., Mirror_1, Mirror_2) and two exemplars of a ‘bottom’ object (e.g., sink_1, sink_2) (see Figure 1A). We combined these top/bottom elements into vertical pairs whose spatial configuration was either *typical* (i.e., top object shown above bottom object) or *atypical* (i.e., top object shown below bottom object). As shown in Figure 2, Contextual-Match pairs were comprised of a top and a bottom object from the same real-world context (e.g., Mirror + Sink); Contextual-Mismatch pairs were comprised of a top and a bottom object from different real-world contexts (e.g., Mirror + Stove). The full combinatorial stimulus set contained 96 contextual-match pairs (48 typical/48 atypical) and 1056 contextual-mismatch pairs (528 typical, 528 atypical). During the experiment, the greyscale object pairs appeared centrally on a uniform grey background, sized at 4×8 cm such that they spanned approximately 3.3×6.5 degrees of visual angle when viewed at a distance of 70 cm.

**Figure 1.**
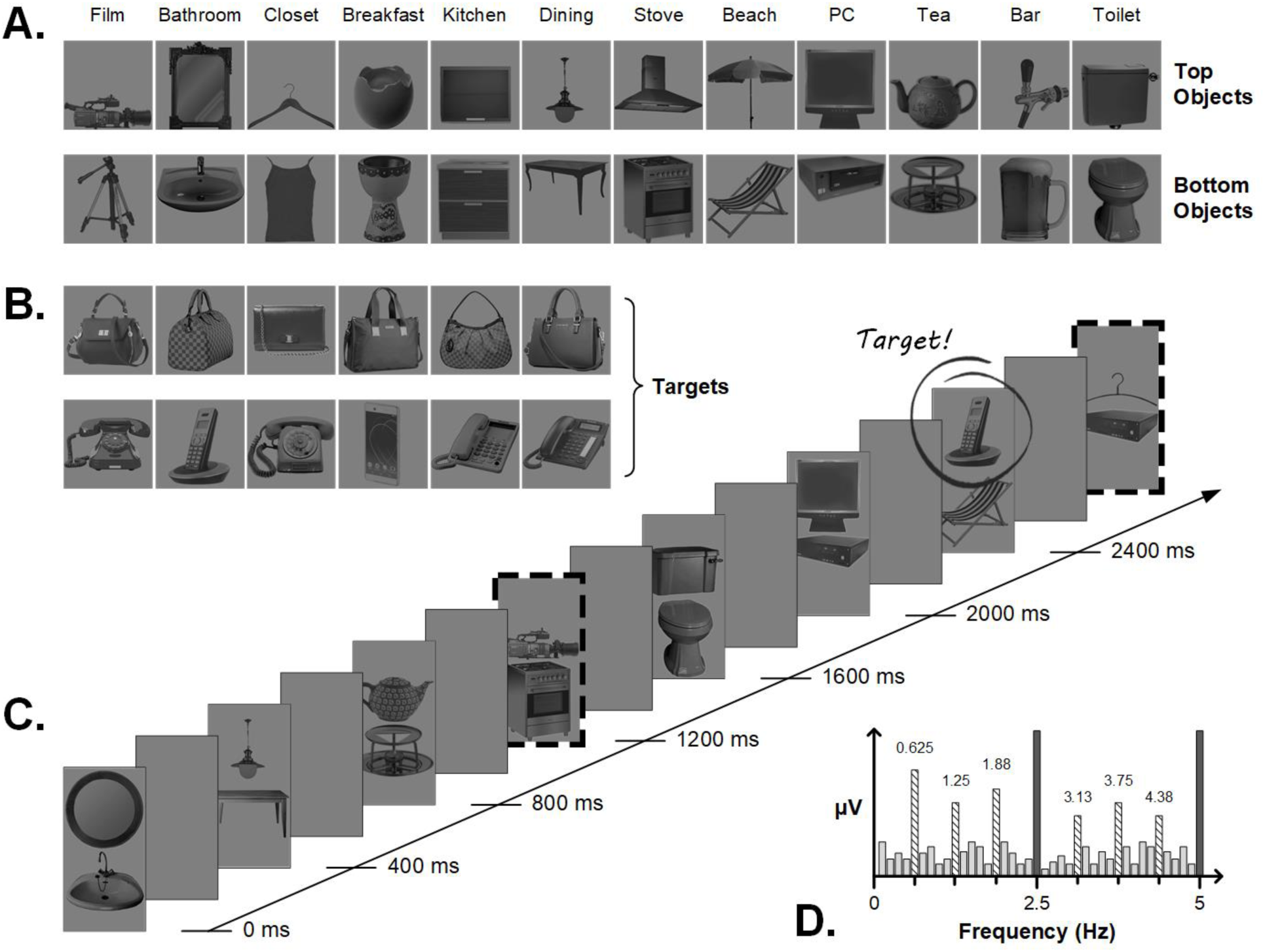
Stimuli and trial sequence details. **A**. Examples of top and bottom objects from the 12 real-world contexts; other exemplars of each object not shown here. Pairing top and bottom objects within- and across-context resulted in contextual-match and contextual-mismatch pairs respectively (e.g., mirror-sink vs. mirror-deckchair). **B**. Target items that participants had to detect during the sequences. **C**. A (truncated) example sequence for the Upright, Typical condition. Object pairs appeared at a rate of 2.5 Hz in the order match, match, match, mismatch, for a mismatch frequency of 0.625 Hz (i.e., 2.5 Hz/4, indicated by dashed borders). Targets could appear randomly in either location at any point during the sequence, except at the mismatch frequency itself. **D**. Toy representation of the expected frequency spectrum: where a response at 2.5 Hz and harmonics reflects general visual processing of all object pairs, a response at the mismatch frequency of 0.625Hz and harmonics captures the differential response to mismatch compared to match pairs.

**Figure 2.**
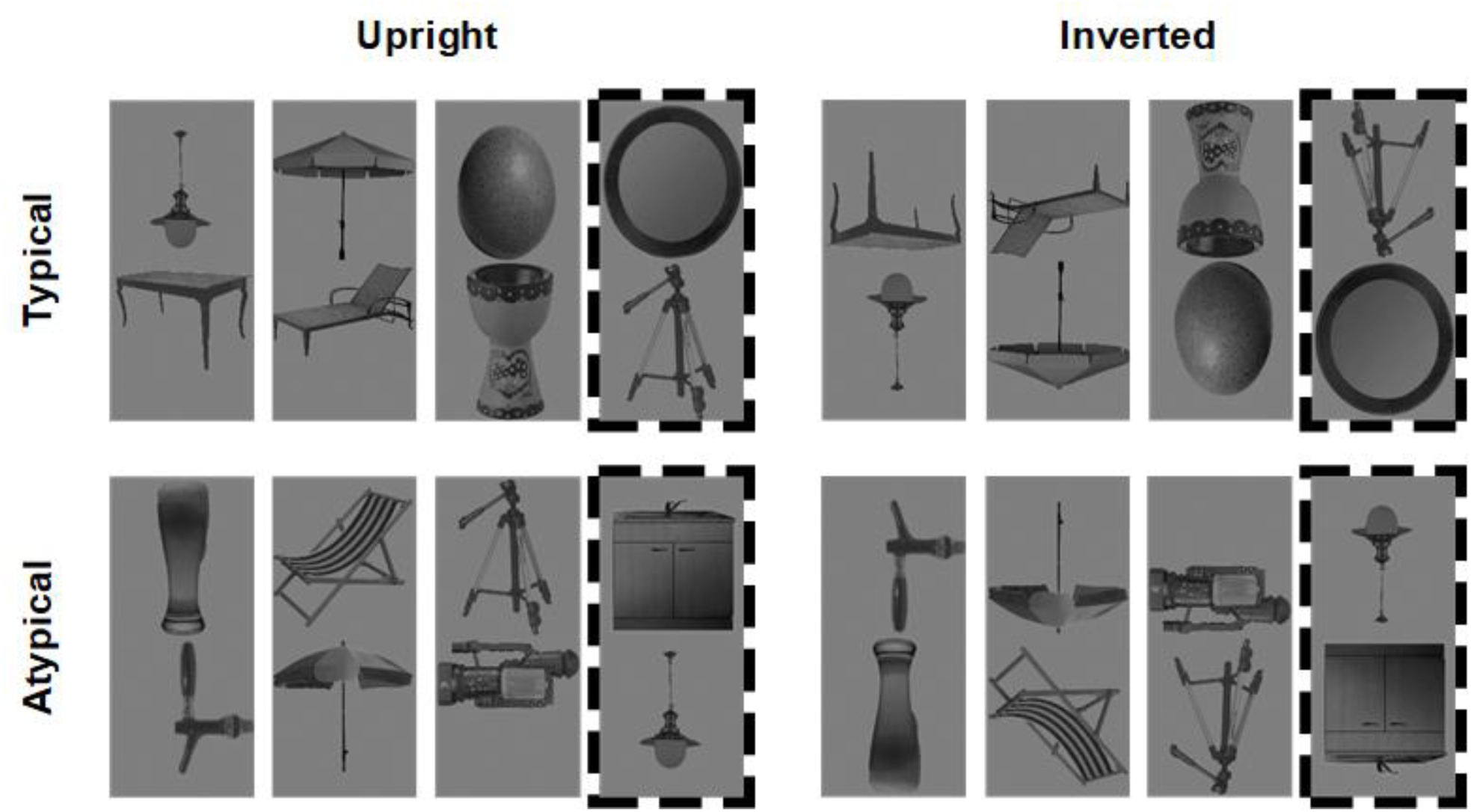
Example contextual-match and contextual-mismatch pairs (dashed border) arranged in typical (top row) and atypical (bottom row) configurations, shown here for the upright (left column) and inverted (right column) conditions.

Target stimuli were six different images of handbags and six different images of telephones (Figure 1B). We chose these specific objects as behavioural response targets since they were not strongly associated with any of the 12 real-world contexts used in the main stimulus set, and therefore no spatial configuration typicality status (i.e., they could not reasonably be considered a ‘top’ or ‘bottom’ object).

### Procedure & Design

The experiment was carried out in a separated, dimly lit testing room that the experimenter visually monitored via webcam from the adjacent room. We used custom software written in Java to display stimuli on a BenQ XL2420Z computer monitor (120Hz refresh rate) fixed at a viewing distance of 70cm. Each presentation sequence lasted 80 seconds and consisted of 200 object pairs from either the typical or atypical stimulus set, presented at a rate of exactly 2.5 Hz (i.e., 400ms SOA, see movie in Supplemental Material). We implemented a squarewave contrast modulation (0-90%) with a 50% duty cycle for each stimulus, such that each object pair appeared onscreen for 200ms, followed by a 200ms blank ISI. Within a sequence, there were 150 contextual-match pairs and 50 contextual-mismatch pairs, such that each of the 48 contextual-match pairs appeared at least 3 times during a single sequence (randomised order; no replacement). Since the full combinatorial set of contextual-mismatch pairs greatly exceeded the number of mismatch stimuli required for each sequence, we divided the 528 mismatch pairs for each typicality condition into 6 sets of 88 pairs, and assigned each set to serve as the stimulus pool for the mismatch stimuli on one sequence. This yielded 6 typical and 6 atypical sequences, in which the 50 mismatch stimuli were randomly drawn without replacement from the relevant set of 88.

Within each sequence, object pairs always appeared in the strict order *match, match, match*, ***mismatch***… (Figure 1C). In this way, our design contained two critical visual periodicities: the image presentation rate (2.5 Hz) and the contextual-mismatch rate (0.625 Hz, i.e., 2.5Hz/4). The embedded periodicity of mismatch pairs is the critical manipulation, as it enables us to quantify the selective response to this stimulus type: here, a neural response at 0.625 Hz reflects the differential response to mismatch pairs amongst match pairs, where aspects of the response that are common to both match and mismatch instances will be captured at 2.5 Hz (visual representation given in Figure 1D). Importantly, since match and mismatch pairs are comprised of the same individual elements, any periodic signal we observe at the mismatch frequency is unlikely to be driven by visual differences between match and mismatch pairs (Figure 2) or by the absolute position of the objects in a stream. Therefore, the response at 0.625 Hz – referred to here as the *Contextual Integration Response –* likely reflects the integration of high-level information about the object identities in each pair. It should be noted that other factors may also contribute to the response at the mismatch frequency: the relatively low number of repetitions of the mismatching pair images, the relatively reduced visual similarity of objects in mismatch pairs, and the reduced probability of targets at the mismatch frequency (see next paragraph). These factors are inherent to the design. Importantly, these factors were equated across the four conditions of interest and should thus not affect the statistical comparisons reported here.

We encouraged participants to attend to the objects in the sequence by engaging them in an orthogonal target detection task in which they pressed the spacebar whenever a handbag or a phone appeared in the stimulus sequence. These targets were not singletons, but randomly replaced either the top or bottom object of a contextual-match pair, such that they always appeared alongside another object (just as in the normal pairs). Per sequence, there were 13 target instances which could appear with equal probability in either position at any moment during the full 86 second stimulation period (but never at the mismatch frequency). Because the targets occurred at random (i.e., nonperiodic) intervals in the sequence, signal arising in response to target instances could not impact the Contextual Integration Response arising at 0.625 Hz. Task performance was monitored online and verbal feedback given; since performance on this orthogonal task was not of interest to the current question, these data were not analysed further.

Since it could be the case that low-level visual differences between the typical and atypical pair arrangements account for differences in the Contextual Integration Response elicited by these conditions, we also included a control condition in which all parameters remained the same, but the entire display was inverted. Inversion preserves the pixel-level differences between the typical and atypical pairs, while attenuating the observer’s ability to process high-level identity information about the objects and their contextual association. This led to a 2×2 design with the factors *Pair Configuration* (typical, atypical) × *Image Rotation* (upright, inverted). Each sequence was preceded and followed by an additional 3 seconds during which the overall contrast of the images ramped up and down respectively (i.e., fade in / fade out periods). There were six trials per design cell for a total of 24 sequences in the full experiment, presented in a pseudorandomised order across two blocks for every participant.

### EEG acquisition

We recorded scalp EEG using a 64-channel active electrode actiCAP system (500 Hz sample rate) with customised electrode positions adapted from the actiCAP 64Ch Standard-2 system (ground electrode placed at AFz; TP10 placed on right mastoid to serve as an additional reference electrode). There were two right frontal channels (AF4, AF8) whose recording was not possible in most participants due to equipment malfunction (i.e., dead electrodes removed from analyses, see pre-processing). Individual electrode offsets were verified to be between 20-50 kOhm before recording commenced. Data were referenced online to the left mastoid and filtered between 0.016-125 Hz using BrainVision Recorder. For all except two participants, we monitored eye movements using external passive electrodes placed at outer canthi of both eyes (horizontal) and immediately above and below the right eye (vertical). These external channels were referenced to a ground electrode placed on the tip of the nose. We synchronised the EEG recording with the start of each stimulation sequence by using a photodiode placed in the upper left corner of the stimulation PC monitor to send triggers via a USB port to the EEG acquisition PC. We visually monitored the EEG trace throughout the experiment, manually initiating each sequence only after it had been free from muscular and ocular artefact for at least 5 seconds. Sequences during which significant movement artefact occurred were immediately repeated.

### Analysis

#### Pre-processing

We carried out EEG data analysis in Matlab 2015b using the LetsWave 6 toolbox (https://github.com/NOCIONS/letswave6), following typically-implemented procedures for this paradigm (Liu-Shuang *et al*. 2014; Rossion *et al*. 2015; Retter and Rossion 2016; Quek and Rossion 2017). We removed the left and right mastoid channels, as well as two right frontal channels (AF4, AF8) whose recording was not possible in most participants due to equipment malfunction (i.e., dead electrodes). For the remaining 61 electrodes, we applied a 4^th^ order Butterworth bandpass filter (0.05-120 Hz) to the raw EEG trace, followed by an FFT multi-notch filter to remove electrical line noise at 50, 100, and 150 Hz. We then segmented each participant’s EEG recording into 24 × 90 s epochs around each stimulation sequence, and quantified the number of eyeblinks within each by inspecting the vertical EOG channel for deflections >100µV. For participants whose mean blinks-per-second exceeded 0.2 (*n* = 12), we used independent component analysis (ICA) with a square mixing matrix to remove the component corresponding to blinks (identified by visual inspection of the topographical distribution and waveform). Next, we visually identified artefact-ridden channels and replaced these using the linear interpolation of the neighbouring 3 channels (maximum 3 channels/participant replaced). Epochs containing a large number of artefact-ridden channels were considered unsuitable for interpolation and removed (an average of 0.65 epochs per participant). We re-referenced each channel’s signal to the average of all 61 scalp channels, before re-cropping each epoch to synchronise with the start of the first full-contrast cycle (i.e., after the fade-in period). This final 78.4 second segmentation corresponded to exactly 196 object-pair presentation cycles, of which 49 were contextual mismatch instances.

#### Frequency Domain

Here we averaged each participant’s final segmentation epochs to obtain *i)* a mean for each condition, and *ii)* a grand mean averaged across all conditions. We subjected the resulting waveforms to Fast Fourier Transformation (FFT) to produce amplitude spectra with a frequency resolution of 0.0128 Hz.

##### Harmonic selection

To identify which harmonics of the image presentation and mismatch frequencies carried significant signal, we first averaged the grand mean amplitude spectra of all 37 participants before pooling the 61 channels to create a group-level scalp-averaged dataset. At each frequency bin in this amplitude spectrum, we calculated a *z*-score by subtracting the mean amplitude of the surrounding bins (defined as the 8 bins, or 0.102 Hz, either side of the bin of interest, excluding the immediately adjacent bin) from that at the bin of interest, and dividing this value by the standard deviation of the surrounding bins. Using a significance criterion of *z* > 1.64 (i.e., *p* < .05, one-tailed^1^), we identified significant signal at the first 19 harmonics of the image presentation frequency (range: 2.5-47.5 Hz), and the first six harmonics of the mismatch frequency, excluding the overlapping frequency at 2.5 Hz (i.e., 0.625, 1.25, 1.875, 3.125, 3.75, & 4.375 Hz).

##### Participant selection

Since our primary goal was to determine whether the Contextual Integration Response was sensitive to the typicality of object configuration, we aimed to restrict our condition-wise analyses to those observers for whom a significant response at the mismatch frequency could be identified when averaging across all conditions and scalp channels. For this condition-blind analysis, we segmented each participant’s grand mean amplitude spectrum around each of the six identified mismatch frequency harmonics (21 bin range, i.e., middle signal bin plus 10 noise bins either side), and summed the values on each bin across the six segments. We then averaged the resulting summed spectra across all 61 scalp channels, and calculated a *z*-score for the middle signal bin using the same procedure and surrounding bin range described above (i.e., 8 bins either side of the middle bin). Participants whose scalp-averaged, condition-blind *z-*score exceeded 1.64 (i.e., *p* < .05, one-tailed) were considered to have a significant Contextual Integration Response; this was the case for 28/37 observers (i.e., 76% of our participant sample; binomial test against chance: *p* < .001). All subsequent tests for the (orthogonal) effects of our experimental manipulations on the Contextual Integration Response were carried out in this observer group.

##### Baseline-correction

For each participant in the identified subset, we repeated the segmentation/summing procedure described above for each of the four conditions separately (note there was no channel averaging here). We then baseline-corrected each participant’s grand and conditional means by subtracting the mean amplitude of the surrounding noise bins from the amplitude at the middle signal bin (as above, we defined the noise range as the 8 bins either side of the middle signal bin, excluding the immediately adjacent bin). The resulting values on the middle signal bin comprised the *Contextual Integration Response* we report below. We carried out statistical analysis of the Contextual Integration Response using repeated-measures ANOVAs with Greenhouse–Geisser corrections applied to degrees of freedom whenever the assumption of sphericity was violated. We used paired *t*-tests for follow-up comparisons (two-tailed, unless otherwise specified), and considered these significant only if they survived Bonferroni correction. Below we report uncorrected *p*-values to enable readers to evaluate the results at the significance criterion of their choosing.

##### Channel inspection

We began by inspecting the condition-wise pattern of means on the channel generating the largest Contextual Integration Response averaging across all conditions (electrode PO10, see Figure 4A inset). We then subsequently expanded our channel selection around this critical electrode to form a larger right hemisphere ROI encompassing multiple occipitotemporal electrode sites. To determine which channels should contribute to this ROI, we averaged the summed grand mean amplitude spectra across our identified subset of observers. For each channel in this group-level spectrum, we calculated a *z* score for the middle signal bin and identified contiguous channels within the right hemisphere where *z* > 4.3 (i.e., *p* < .00001, one-tailed). To support our subsequent enquiry into the lateralised nature of the Contextual Integration Response, we implemented the same channel selection procedure within the left hemisphere.

**Figure 3.**
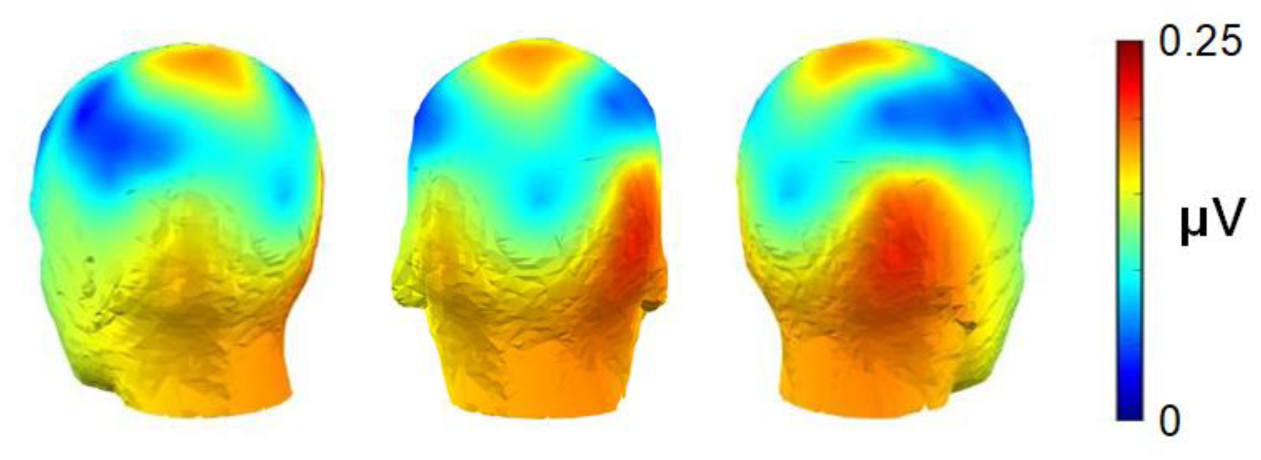
The group-level periodic response elicited by contextual-mismatch pairs, averaged across all conditions. This Contextual Integration Response was strongest over right occipitotemporal electrode sites, with maximal activation on electrode PO10.

**Figure 4.**
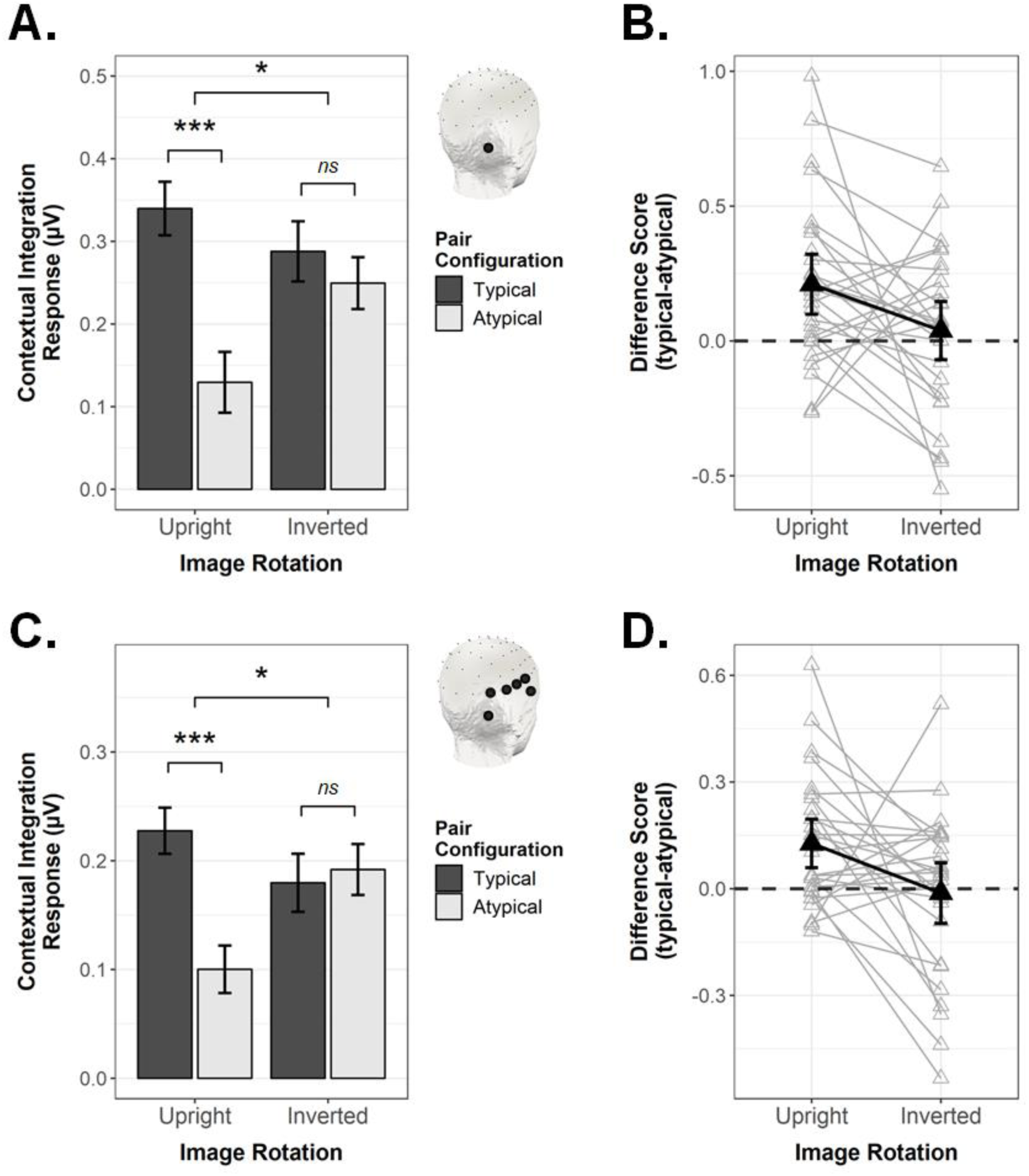
The Contextual Integration Response as quantified in the frequency domain, shown as a function of Pair Configuration and Image Rotation. **A**. Conditional mean response magnitude on electrode PO10 (see inset) Error bars are within-subjects SEM; \**p* < .05, \*\**p* < .01, \*\*\**p* ≤ .001. **B**. Difference scores (typical – atypical) for electrode PO10, shown with 95% confidence intervals (CIs). Black triangles are group means; grey triangles are individual participant means. **C**. & **D**., as above, as averaged across the right occipitotemporal ROI.

#### Time Domain

We isolated the neural signal evoked by contextual-mismatch pairs (i.e., the mismatch-selective response) in the time domain by applying a 4^th^ order Butterworth low-pass filter (30 Hz cut off) to each participant’s final segmentation epochs, followed by an FFT multi-notch filter to remove signal carried at the first 19 harmonics of the image presentation rate (range: 2.5-47.5 Hz; slope cutoff width = 0.05). Next we segmented a 650ms epoch around each contextual-mismatch instance (−200 to 450ms), resulting in 48 epochs per trial sequence. For each participant, we averaged these epochs within condition, and baseline-corrected the resulting waveforms by subtracting the mean amplitude of period from −200ms to 0ms from the amplitude at every timepoint. Group-level scalp topographies for each condition were obtained at 50ms time increments following mismatch onset.

## Results

### Frequency Domain

Figure 3 shows the spatial distribution of the Contextual Integration Response averaged across all conditions. At the group-level, this signal was strongest over right occipitotemporal electrode sites, with the largest condition-averaged response evident on electrode PO10, which was the highest responding channel for 9/28 included participants.

As can be seen in Figure 4A, inspecting the signal on electrode PO10 as a function of condition revealed that presenting object pairs in their typical real-world configurations enhanced the Contextual Integration Response for upright stimuli, *t*(27) = 3.70, *p* < .001, *d* = 0.70. Crucially, however, this effect was specific to upright displays: For inverted displays, there was no influence of *Pair Configuration* on the magnitude of the Contextual Integration Response, *t*(27) = 0.70, *p* = .492, *d* = 0.13. The interaction between *Pair Configuration* and *Image Rotation* was significant in a 2-way repeated measures ANOVA, *F*(1,27) = 4.63, *p* = .041, *η*^2^G = 0.027.

Figure 3 shows that the Contextual Integration Response was not solely evident on electrode PO10, but that a broader right occipitotemporal region carried this signal of interest. Thus, we next considered whether the interactive pattern observed for PO10 would be maintained when considering the response averaged across electrodes PO10, PO8, P8, TP8, T8, FT10 (see Methods for details on how we selected these channels). Averaging across these right OT channels (Figure 4C & 4D), we observed the same interactive pattern as was found at single electrode level, *F*(1,27) = 5.83, *p* = .023, *η*^2^G = 0.047. Where *Pair Configuration* plainly modulated the Contextual Integration Response for upright displays, *t*(27) = 3.66, *p* = .001, *d* = 0.69, it exerted no influence on the magnitude of the response when the display was inverted, *t*(27) = −0.28, *p* = 0.781, *d* = 0.05.

Lastly, we asked whether the interactive pattern demonstrated for right OT channels could be considered a truly lateralised effect. Using the same channel selection procedure as implemented for the right ROI (see Methods), we averaged the Contextual Integration Response across four contiguous occipitotemporal electrodes within the left hemisphere (PO9, PO7, P7, FT9). We then conducted a 3-way repeated-measures ANOVA with the goal of determining whether the interaction between *Image Rotation* and *Pair Configuration* would be further qualified by *ROI* (left, right). This was indeed the case, *F*(1,27) = 4.95, *p* = .035, *η*^2^G = 0.010, leading us to conduct a follow-up two-way repeated-measures ANOVA for the left ROI (right ROI already reported above). In the left ROI (Figure 5A, left column), we observed that *Image Rotation* and *Pair Configuration* did not interact significantly, *F*(1,27) = 0.11, *p* = .744, *η*^2^G = 0.001, although there was a significant main effect of *Pair Configuration, F*(1,27) = 8.86, *p* = .006, *η*^2^G = 0.082 (main effect of *Image Rotation* also not significant, *F*(1,27) = 3.09, *p* = .090, *η*^2^G = 0.020). To verify that the key 3-way interaction did not simply arise due to a different number of channels contributing to the left and right ROIs, we repeated the 3-way ANOVA after restricting the right ROI channels to mirror those on the left (i.e., average of PO10, PO8, P8, FT10 *vs*. average of PO9, PO7, P7, FT9, see insets in Figure 5A). This did not change the pattern of effects in any way, that is, the 3-way interaction remained reliable, *F*(1,27) = 5.07, *p* = .033, *η*^2^G = 0.011. A follow-up ANOVA for the (reduced) right ROI (Figure 5A, right column) showed a significant interaction between *Image Rotation* and *Pair Configuration, F*(1,27) = 5.36, *p* = .028, *η*^2^G = 0.044, which paired *t*-tests indicated was due to a significant *Pair Configuration* effect for the upright, *t*(27) = 3.81, *p* < .001, *d* = 0.72, but not the inverted condition, *t*(27) = 0.17, *p* = .868, *d* = 0.03.

**Figure 5.**
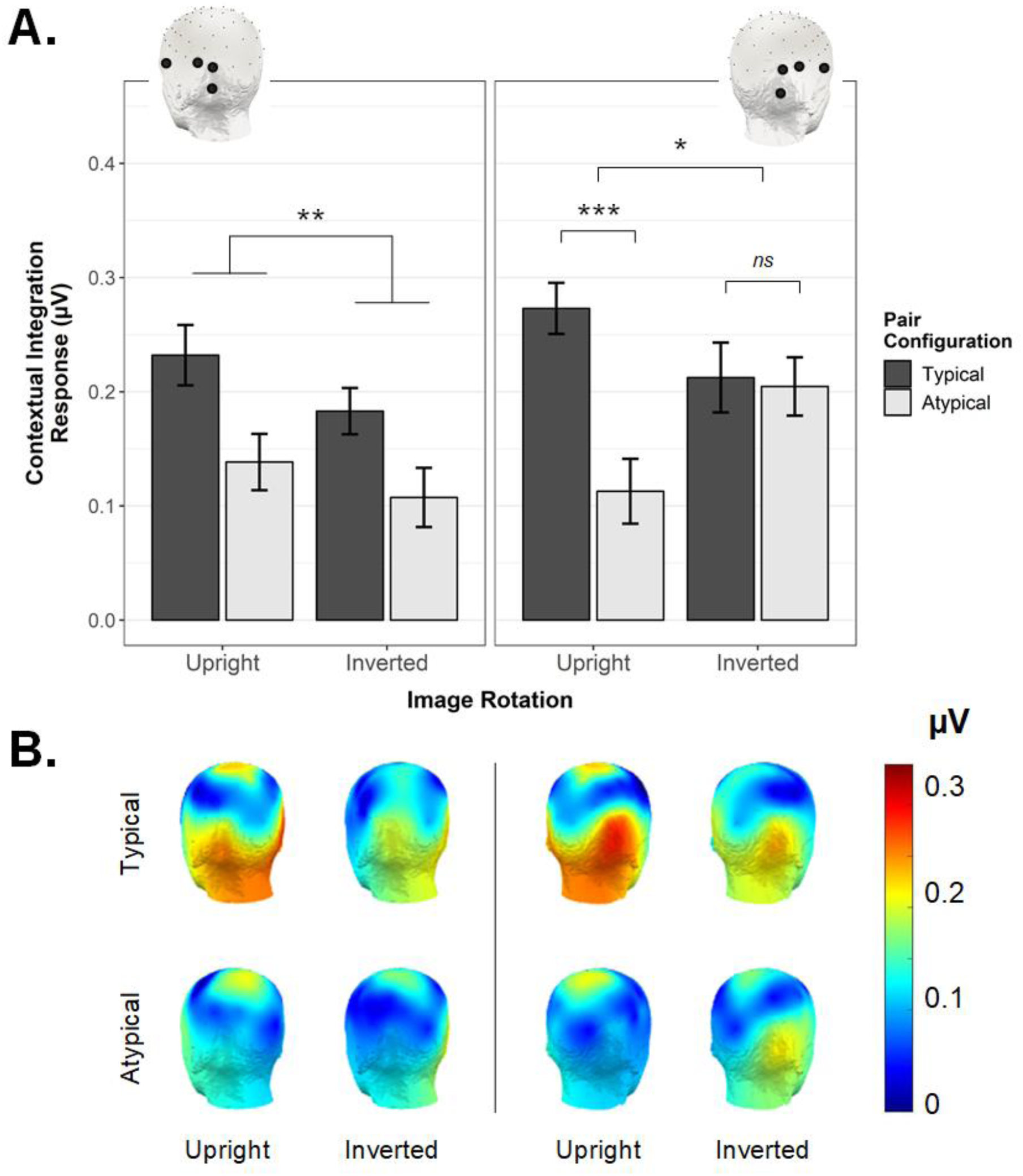
**A**. The significant 3-way interaction between *Pair Configuration, Image Rotation*, and *ROI* observed using identical left and right OT ROIs (insets). In the right ROI, the Contextual Integration Response was stronger for typical compared to atypical pairs (only when the display was upright, but not inverted). In the left ROI, only a main effect of Pair Configuration was evident. Error bars are within-subjects SEM; *p < .05, **p < .01, ***p < .005. **B**. Scalp topographies corresponding to the Contextual Integration Response observed in each condition (posterior left and right views).

### Time Domain

Where the above analyses establish the relative importance of the right occipitotemporal region in driving the critical pattern of interest, this frequency-based approach does not capture information about either the polarity or timecourse of mismatch-selective activation over any particular electrode site. As such, we next inspected the temporal unfolding of the Contextual Integration Response as a function of time from mismatch onset. Figure 6B shows the grand mean mismatch-selective waveform averaged across both ROIs (corresponding scalp topographies given in Figure 6A). This representation of activation over time is the direct analogue of the Contextual Integration Response as quantified in the frequency domain (see Figure 3), since here the response at the image presentation frequency (i.e., 2.5 Hz and harmonics) has been removed with notch-filtering (see Methods). As highlighted in Figure 6B, local peaks in the grand mean Contextual Integration Response were evident at 130ms and 318ms.

**Figure 6.**
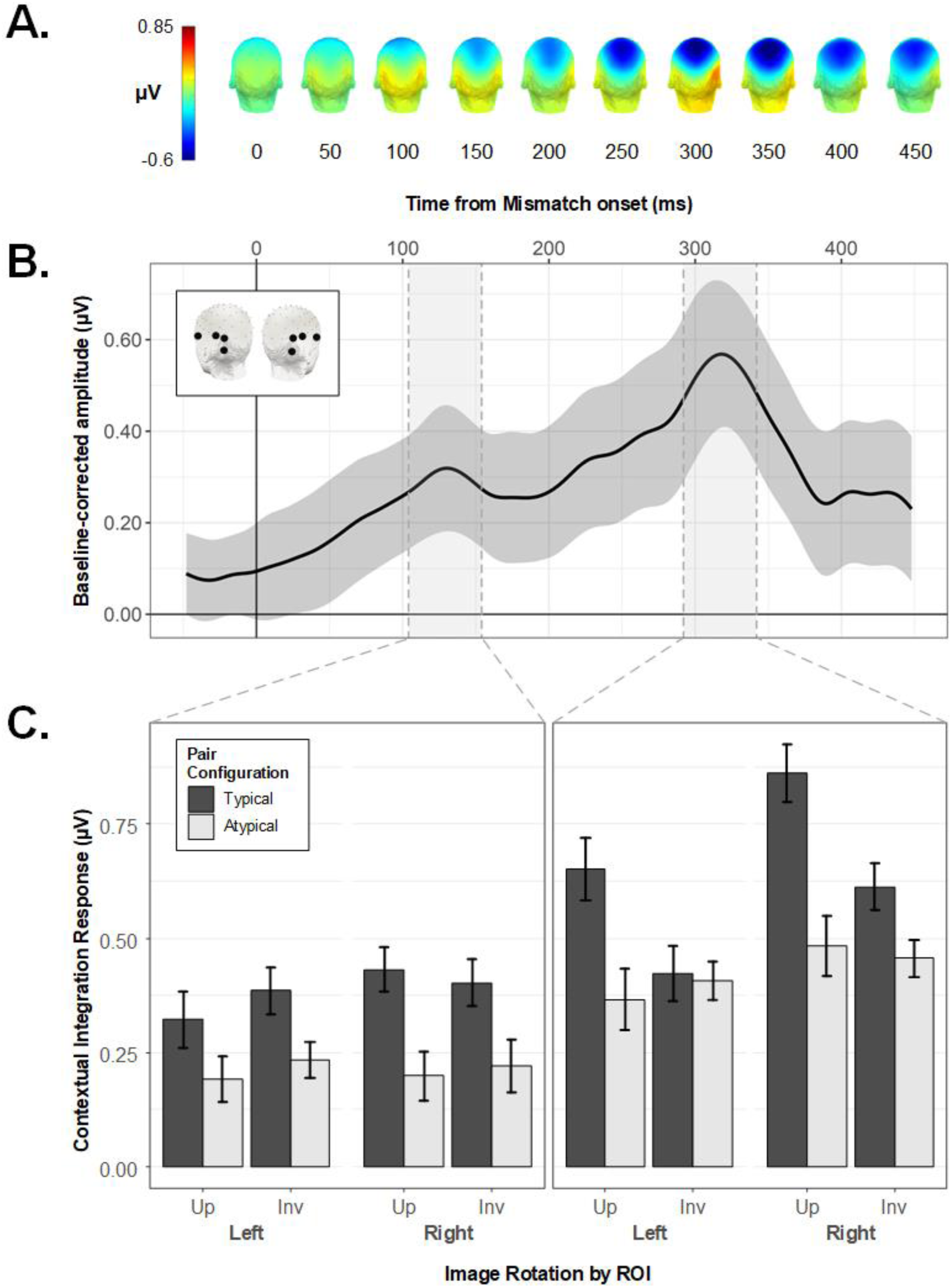
The Contextual Integration Response in the time domain. **A**. Scalp topographies for the grand averaged mismatch-selective response, shown at 50ms intervals following mismatch onset. **B**. The grand-averaged mismatch-selective waveform with 95% confidence band, averaged across symmetrical left and right OT ROIs (see inset). Shaded regions reflect 50ms time windows around identified peaks at 130ms and 318ms. **C**. Conditional mean peak amplitudes within symmetrical left and right ROIs. Where amplitude at 130ms was larger for Typical *vs*. Atypical pairs regardless of image orientation (left column), by 318ms this effect of *Pair Configuration* was only evident for Upright object pairs, and not Inverted ones (right column). Error bars are within-subjects SEM.

We isolated the identified peaks for analysis by averaging the amplitudes in each condition across a 50ms window centred on each latency (see Figures 7B and 6C) and subjecting the resulting values for each peak to a repeated-measures ANOVA with the factors *ROI, Image Rotation*, and *Pair Configuration*. Figure 6C shows the conditional mean amplitudes for the 130ms and 318ms peaks. For the former, only the main effect of *Pair Configuration* reached significance, *F*(1,27) = 12.11, *p* = .002, *η*^2^G = 0.075, with higher peak amplitudes in the typical (*M* = 0.39, *SEM*_*within*_ = 0.03) *vs*. atypical condition (*M* = 0.21, *SEM*_*within*_ = 0.03). In contrast, the later peak was characterised by a significant interaction between *Pair Configuration* and *Image Rotation, F*(1,27) = 7.48, *p* = .011, *η*^2^G = 0.026. Unlike the frequency domain analysis, here the critical interaction between these factors was not qualified by *ROI, F*(1,27) = 0.19, *p* = .667, *η*^2^G = 0.0002. Follow-up paired *t*-tests on this later time window indicated that for upright sequences, peak amplitudes were higher in the context of typically configured (*M* = 0.76, *SEM*_*within*_ = 0.05) *vs*. atypically configured pairs (*M* = 0.42, *SEM*_*within*_ = 0.05), *t*(27) = 3.72, *p* < .001, *d* = 0.71. For inverted object sequences, there was no difference between typical (*M* = 0.52, *SEM*_*within*_ = 0.04) and atypical (*M* = 0.43, *SEM*_*within*_ = 0.04) peak amplitudes, *t*(27) = 1.44, *p* = .161, *d* = 0.27. The same analysis implemented with a wider time window of 100ms around each peak latency produced the same pattern of effects (Main effect of *Pair Configuration* at 130ms, *F*(1,27) = 8.60, *p* = .007, *η*^2^G = 0.060; *Pair Configuration* × *Image Rotation* interaction at 318ms, *F*(1,27) = 4.99, *p* = .034, *η*^2^G = 0.023).

**Figure 7.**
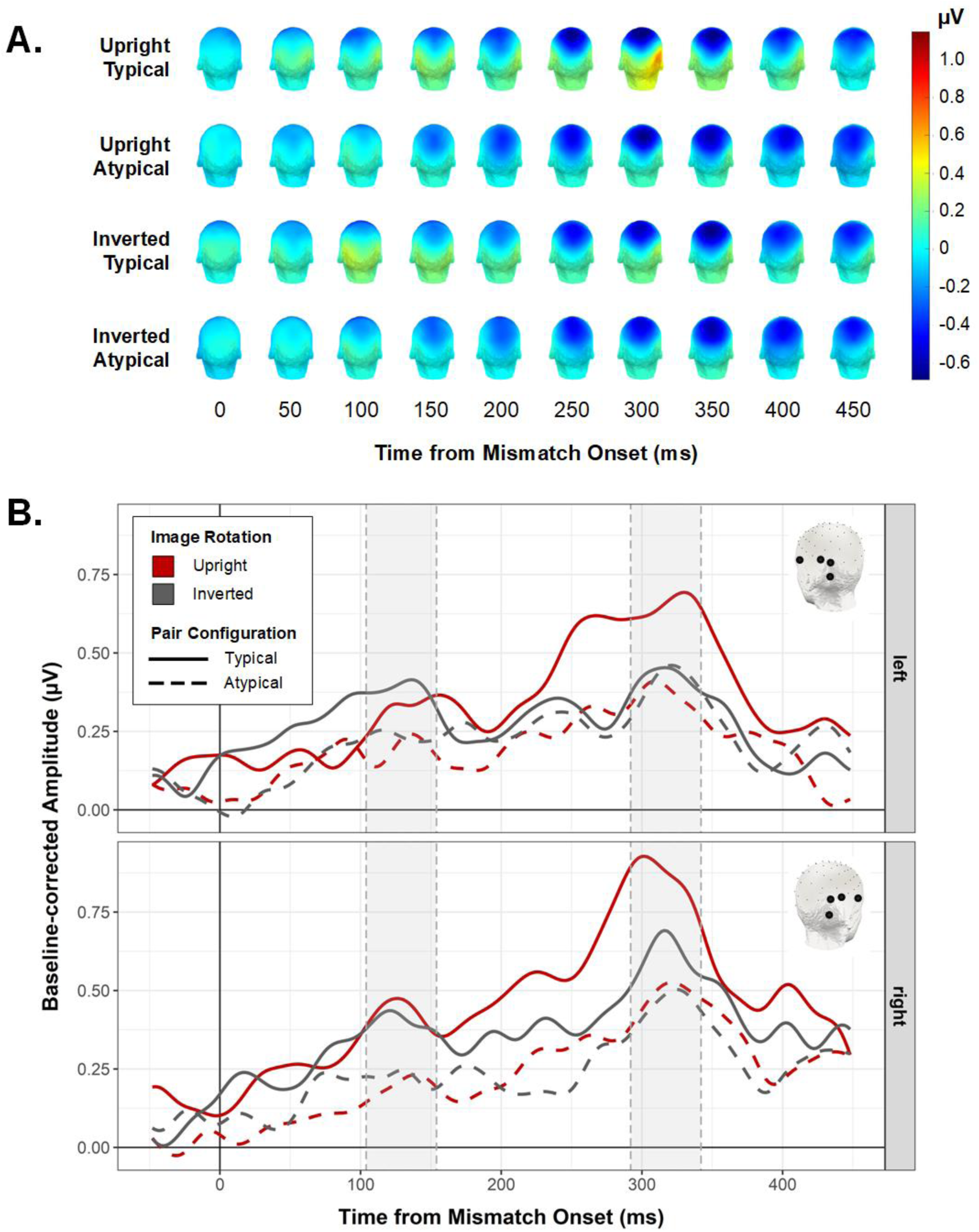
**A**. Scalp topographies for each condition, shown at 50ms intervals following mismatch onset. **B**. Mismatch-selective waveforms for condition in symmetrical left (top row) and right (bottom row) ROIs. Shaded regions indicate peak windows subjected to statistical analysis (see Figure 6C).

## Discussion

The current study focused on a neural signal over occipitotemporal electrode sites that reflected the degree to which responses evoked by contextually non-associated objects differed from those evoked by contextually-associated objects. Since these so-called match and mismatch object pairs were comprised of exactly the same object elements (combined either within or across real-world contexts), we treated this Contextual Integration Response as a direct index of the neural representation of identity-based associations between simultaneously presented objects (i.e., whether or not they belong to the same real-world context). The key finding of our study is that this identity-association index, present in 28/37 participants, is facilitated by presenting objects in their typical real-world relative positions (e.g., mirror above sink *vs*. mirror below sink). Since presenting objects in their typical configurations enhanced the degree to which their identity-based association was represented in the brain, this suggests that these two high-level dimensions along which objects in natural scenes relate to each other influence object processing interactively. Importantly, the data presented here rule out a low-level explanation for this modulation. For example, it could be the case that typical objects configurations have better overall contour alignment than atypical ones, thereby benefiting from increased Gestalt-like grouping. Inversion provides an ideal way to exclude such low-level contributions, as it hampers access to high-level information about object identity and inter-object relations, while leaving image-level differences between typical and atypical pair configurations perfectly intact. In our data, inverting the object displays eliminated the effect of positional regularity as quantified in the frequency domain. Additional time domain analysis revealed this high-level effect of real world positioning manifested in the Contextual Integration Response during a peak approximately 320ms post stimulus onset, but not during an earlier peak around 130ms.

### Identity and location-based associations are bound at the level of object processing

In the current study, both identity- and position-based associations between objects were orthogonal to the observer’s task, which was to detect exemplars belonging to two different target object categories (handbags and telephones). Since these target categories had no contextual association with the other object stimuli, and since targets appeared with equal probability in both the top and bottom object locations, attending to the *relationship* between concurrent objects (either their contextual association or positional regularity) did not serve target detection performance in any way. Thus, task strategies (e.g., holding multiple shape templates in mind) were not related to our manipulation of inter-object relations. As such, our finding that presenting objects in their typical real-world relative positions enhances the representation of their identity-based association suggests that contextual associations between objects are tightly linked to their typical relative positions, with these regularities jointly facilitating object perceptibility *itself* (Biederman *et al*. 1982; Bar 2004; Kaiser *et al*. 2019).

The extent to which identity- and position-based inter-object associations interactively modulate object processing must depend on the relative strength of these associations, which need not necessarily co-vary (e.g., a pencil and eraser have a very strong contextual association, but no corresponding typical spatial configuration). Interactive effects of identity- and position-based associations are presumably strongest for stimuli characterised by strong regularities along both dimensions – that is, real-world object pairs for which it would be unusual to encounter one object without the other, and never in the atypical configuration (as we used here). The present results speak to the interactive influence of identity- and position-based associations on the processing of such coherent and familiar object groups, i.e., wherein the individual object elements arguably comprise parts of a unified conceptual whole (e.g., cistern above toilet, see Figure 1A), analogous to parts making up an object.

The data here also shed light on the timecourse over which high-level associations between objects might interact. Specifically, we found that presenting objects in their typical relative positions enhanced the strength of the Contextual Integration signal over occipitotemporal electrode sites during both an early (∼130ms) and later peak (∼320ms). Importantly, however, where the earlier peak manifested this effect for both upright and inverted displays, during the later peak this facilitation by typical object arrangements was evident only for upright object pairs. We interpret this as the earlier stages of the Contextual Integration Response being sensitive to perceptual grouping effects (i.e., grouping driven by low-level visual properties, such as enhanced contour alignment for Typical *vs*. Atypical arrangements), with the high-level influence of spatial associations on object integration arising comparatively later. In future, time-resolved decoding approaches could be used to more precisely characterise the onset and duration of these low- and high-level grouping effects, which could well have overlapping temporal profiles.

As a related point, it is worth considering whether the contextual integration response peak we observe at ∼320ms is similar in kind to the ERP differences in similar time windows evoked by objects that violate semantic expectations induced by scene primes (Ganis and Kutas 2003; Goto et al. 2009; Võ and Wolfe 2013). While contextual mismatch presumably underpins the responses in both our own study and these previous ones, we believe that they may not be directly comparable, for two reasons. First, for scene-object (in)consistencies, ERPs to objects violating scene-induced expectations do not seem to be modulated by the object’s location in the scene; that is, semantic and spatial violations reportedly do not interact (Võ and Wolfe 2013). By contrast, we observed that the neural response to the contextual association between concurrent objects was highly sensitive to whether or not the objects were arranged in their typical relative positions. As alluded to above, this suggests that the strong interdependence of contextual and positional associations, as observed in the current EEG response over occipitotemporal cortex, may be specific to cases where elements form a unitary conceptual whole. This is unlikely to be the case for scene-object combinations, where a contextually (in)consistent object comprises just one element of a cluttered visual scene, and therefore may not have a strong spatial association with any one other scene element.

Second, previous scene-object studies measured violations of expectations induced by a scene preview, shown several hundred milliseconds before the object’s onset. In these designs, responses evoked by semantically inconsistent objects are thought to reflect (domain-general) violations of semantic expectations, similar to those observed for semantic violations in sentences (Kutas and Federmeier 2011; Võ and Wolfe 2013). Whether or not scene-object inconsistencies evoke similar early (∼300 ms) ERPs when scene and object are presented simultaneously is less well established (Demiral et al. 2012; Mudrik et al. 2014; Lauer et al. 2018), and has yet to be convincingly dissociated from low-level accounts as these studies have lacked an inversion control condition. By contrast, the contextual and positional consistency effects observed in the current study were elicited by brief, simultaneous object displays, in the context of a challenging orthogonal task. The fact that we find evidence of an interaction between contextual and positional information under such conditions suggests that these high-level associations between objects can affect visual processing online, rather than only through prior expectations.

### Where are identity-based and location-based associations bound in the brain?

The spatial distribution of the Contextual Integration Response as reflected in both the frequency (Figure 5B) and time domain (Figures 7B) highlights the relevance of occipitotemporal regions in processing the contextual and positional associations between objects. What might be the neural source of this conjoint representation of identity-based and positional associations between objects? Using fMRI and sequential prime and target presentation, Gronau and colleagues (2008) reported several brain regions showing an interaction between the contextual association and spatial configuration of objects, including inferior prefrontal cortex (IPC), parahippocampal cortex (PHC), and lateral occipital complex (LOC). Other studies have similarly implicated LOC in representing the spatial configuration of objects (Roberts and Humphreys 2010; Kim and Biederman 2011; Kaiser and Peelen 2018). For example, multivariate fMRI response patterns in anterior LOC evoked by regularly (vs. irregularly) positioned object pairs were less accurately modelled by a linear combination of response patterns evoked by the pairs’ constituent objects, suggesting that individual objects were integrated (or grouped) when presented in their typical spatial configuration (Kaiser and Peelen 2018). The current EEG results may similarly reflect grouping for correctly-positioned objects with a strong contextual association, which we argue above could be considered to form a conceptual ‘whole’ (e.g., egg + eggcup). Specifically, contextually-mismatched objects that have no familiar identity-based association (therefore no meaningful spatial association either), may stand out best in the context of an image stream that is predominantly comprised of coherent, recognisable groups.

An intriguing possibility raised by the current results is that the high-level modulatory influence of spatial configuration on the representation of identity associations may be stronger in right visual areas compared to left. In our data, the frequency-based quantification of the Contextual Integration Response demonstrated this effect exclusively over right occipitotemporal electrode sites. In corresponding left occipitotemporal sites, typically-configured pairs elicited a stronger Contextual Integration Response than atypically-positioned pairs for both upright *and* inverted displays, such that we could not exclude the possibility that the effect in this region was driven by low-level visual features. However, subsequent decomposition in the time domain revealed a comparable influence of spatial configuration on contextual object integration in both left and right occipitotemporal sites at approximately 300ms post stimulus onset. For now, the degree to which the high-level interaction between identity and position-based associations between objects should be considered right-lateralised remains unclear, particularly since several existing studies have in fact implicated left hemisphere regions in processing inter-object relations (Gronau *et al*. 2008; Roberts and Humphreys 2010). Whether these differences in lateralisation pattern are meaningfully related to design (e.g., sequential presentation format; Gronau *et al*. 2008), stimuli (e.g. inclusion of tools; Roberts and Humphreys 2010), or task parameters will be an important avenue for future investigation – for now, they underscore the need for future fMRI studies to explicitly consider how identity-based and positional associations between objects interactively facilitate object processing in visual cortex.

### Multi-object grouping in the context of temporal competition

Up until now we have only considered object position within a relative framework (i.e., lamp above table). However, there is increasing evidence that presenting an object in its typical *absolute* location (e.g., lamp in the upper visual field) can also facilitate its visual processing (Quek and Finkbeiner 2014; Kaiser and Cichy 2018a; 2018b; Kaiser et al. 2019). In our design, the Contextual Integration Response could not reflect absolute object position of the individual objects because all objects in a stream were either presented in typical or atypical absolute position, such that absolute positional typicality was not specific to the oddball frequency. However, typical absolute position of the objects could have contributed indirectly by facilitating object integration. Specifically, if object processing is facilitated by presenting the objects in appropriate absolute locations (e.g., lamp in the upper visual field), this may in turn enhance the subsequent integration of object-identities contained in each pair. We think this is unlikely to be the case: First, whereas all objects in our stimulus set are readily identifiable as a top or bottom with respect to their paired object (see Figure 1A), not every object can be associated with an absolute vertical location (e.g., a lamp belongs above a table just as an egg belongs above an eggcup, but only the former appears with statistical regularity in the upper visual field). Confirming the weak absolute positional associations of our stimuli, a previous study using the same stimulus set provided evidence for enhanced detectability of the objects presented in their typical relative positions, but found no effect of typical absolute position (Stein et al. 2015), even though the same detection paradigm was sensitive to such effects for objects that are more strongly associated with particular locations (e.g. airplane, shoe) and presented at larger eccentricities (Kaiser and Cichy 2018b). For these reasons, we think the contribution of typical absolute position is likely to be small in our study, with the objects’ *relative* positioning being the driving factor underlying our results.

Why should the differential response to non-associated objects among contextually-associated objects be larger when objects appear in their typical real-world relative positions? As alluded to above, we believe that this effect may reflect a ‘grouping’ adaptation in the human visual system designed to overcome cortical resource limitations. A wealth of evidence shows that when multiple entities must be processed simultaneously, stimuli compete for overlapping resources, such that the response to each individual stimulus is attenuated. An efficient way to reduce the number of individual items competing for representation may be to *group* highly predictable constellations of objects (e.g., lamp above table; Kaiser *et al*. 2019). Such multi-object grouping effects have previously been demonstrated in the context of spatial competition, using the same, highly unitary object pairs as used here: target detection performance improved when surrounding distractor object pairs were presented in their typical real-world relative positions (Kaiser *et al*. 2014). Observers in our study did not encounter a large number of objects in this type of single, cluttered display, but instead had to contend with a new pair of objects appearing every 400ms in a continuous, long-lasting stream (see movie in Supplemental material). That presenting objects in their typical relative positions enhanced the representation of their contextual association under these conditions sheds new light on visual adaptations for simplifying scene analysis, showing that multi-object grouping based on positional regularities serves to reduce competition between high-level stimuli also when processing is temporally constrained. In the context of such temporal competition, the visual system may be better able to group contextually-associated objects when these objects appear in their typical relative positions, leaving more cortical resources available to ‘notice’ the non-associated object pair than when the objects are atypically-configured, and hence less groupable.

## Conclusions

The results reported here establish that arranging objects with respect to their typical real-world relative positions enhances the degree to which their contextual association is represented in the brain. Our data suggest that identity-based and positional associations between objects do not exert isolated influences on how we perceive and represent objects, but that these high-level dimensions along which objects in natural scenes relate to each other interactively facilitate object processing. We interpret this finding as reflecting the perceptual grouping of co-occurring objects with highly stereotyped relative positions.

## Funding

This work was supported by funding from the European Union’s Horizon 2020 research and innovation programme under the Marie Sklodowska-Curie grant agreement No 841909 (G.L.Q.) and the European Research Council grant agreement No 725970 (M.V.P.).

## Acknowledgements

The authors would like to acknowledge A. Theodorou, L. Billen, N. Parrella, and M. Picó Cabiró for their assistance with data collection.

## Footnotes

1 We use a one-tailed test as here the goal is to identify frequencies at which the signal is significantly *greater* than the noise in the surrounding bins. Instances in which the signal is significantly lower than the surrounding bins are not relevant.

## Notes

### Competing Interest Statement

The authors have declared no competing interest.

### Summary of Updates

Discussion expanded.

